# Bioinformatic and computational analysis reveals the prevalence and nature of PY motif-mediated protein-protein interactions in the Nedd4 family of ubiquitin ligases

**DOI:** 10.1101/2020.11.12.380584

**Authors:** A. Katherine Hatstat, Michael D. Pupi, Dewey G. McCafferty

## Abstract

The Nedd4 family contains several structurally related but functionally distinct HECT-type ubiquitin ligases. The members of the Nedd4 family are known to recognize substrates through their multiple WW domains, which recognize PY motifs (PPxY, LPxY) or phospho-threonine or phospho-serine residues. To better understand substrate specificity across the Nedd4 family, we report the development and implementation of a python-based tool, PxYFinder, to identify PY motifs in the primary sequences of previously identified interactors of Nedd4 and related ligases. Using PxYFinder, we find that, on average, half of Nedd4 family interactions are PY-motif mediated. Further, we find that PPxY motifs are more prevalent than LPxY motifs and are more likely to occur in proline-rich regions. Further, PPxY regions are more disordered on average relative to LPxY-containing regions. Informed by consensus sequences for PY motifs across the Nedd4 interactome, we rationally designed a peptide library and employed a computational screen, revealing sequence- and biomolecular interaction-dependent determinants of WW-domain/PY-motif interactions. Cumulatively, our efforts provide a new bioinformatic tool and expand our understanding of sequence and structural factors that contribute to PY-motif mediated substrate recognition across the Nedd4 family.

## Introduction

Neuronal precursor cell-expressed developmentally downregulated 4 (Nedd4) is the founding member of a family of HECT-type E3 ubiquitin ligases that share a common architecture but have distinct cellular functions. The Nedd4 family is characterized by a multi-domain architecture comprised, from N- to C-terminus respectively, of a C2 domain for membrane localization, two to four WW domains for substrate recognition, and a catalytic HECT domain (**Figure 1A**).^1–5^ As the final enzymes in the ubiquitin signaling cascade, the Nedd4 family of HECT-type E3 ubiquitin ligases receives ubiquitin from a ubiquitin-E2 conjugating enzyme thioester adduct. The ubiquitin-HECT E3 conjugate then passes ubiquitin to a substrate protein via isopeptide bond formation at target lysine residues. Nedd4 and related HECT-type ligases are thus responsible for conferring substrate specificity in the ubiquitin signaling pathway. Understanding the specificity of the Nedd4 family of ubiquitin ligases is of particular interest due to the role of Nedd4 in the regulation of proteostasis in various conditions including cancers^6–8^ and neurodegenerative disorders^9–16^ and with recent insights into the potential of Nedd4 to serve as a therapeutic target.^17–25^

**Figure 1.**
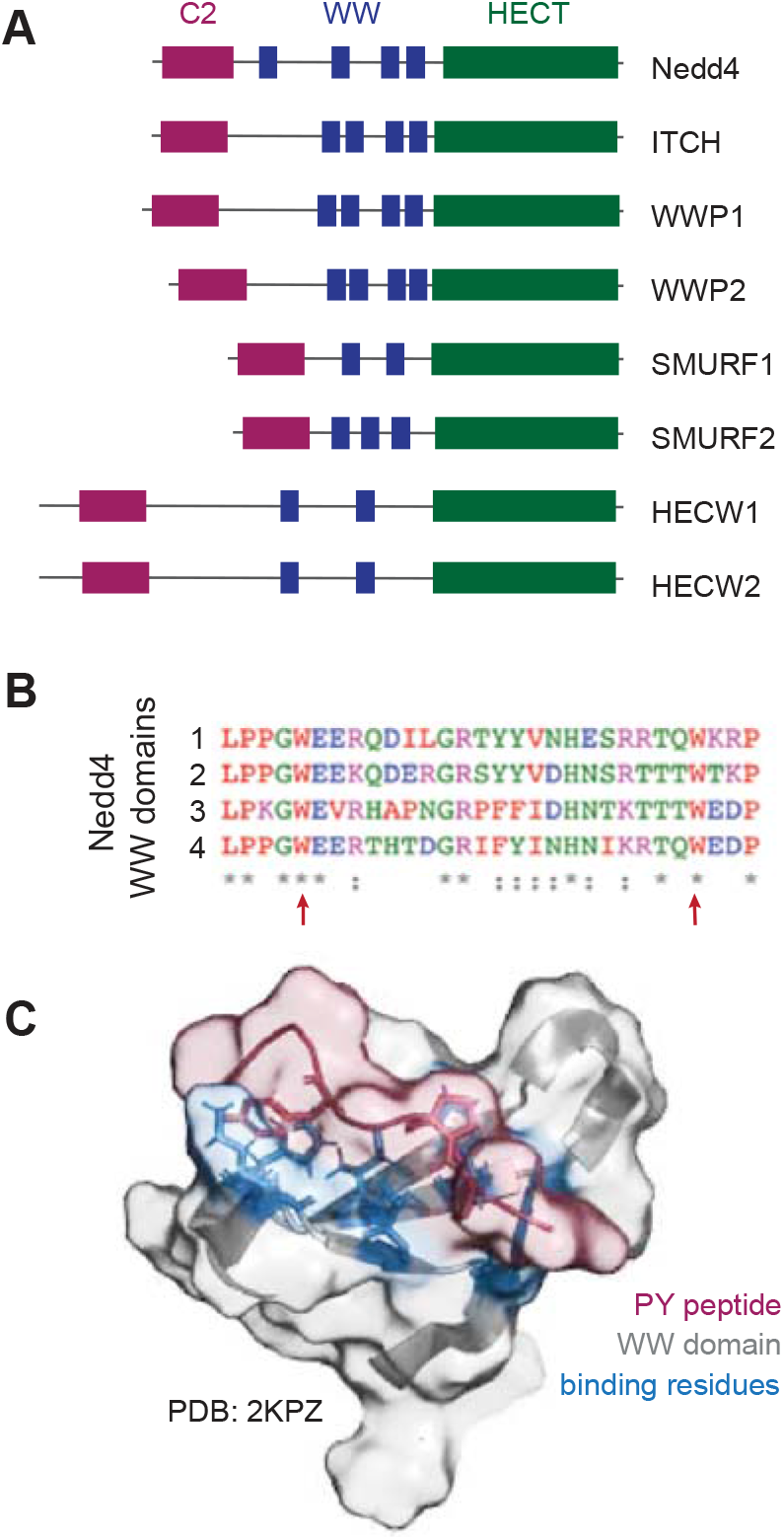
The Nedd4 family of ligases contains conserved WW domains for substrate recognition. **(A)** Representative structure motif diagrams of the Nedd4 family. Nedd4 and related ligases are structurally conserved, containing 2–4 WW domains that recognize substrates containing a PY motif (PPxY, LPxY) or phosphorylated threonine or serine residues. **(B)** Alignment of the four WW domains from prototypical member Nedd4 shows moderate sequence similarity and highlights conserved residues, including the two characteristic tryptophan residues (indicated by red arrows). **(C)** Solution state NMR structure of the Nedd4 WW domain 3 (grey) in complex with a PY motif peptide (red) from a known Nedd4 substrate (PDB ID: 2KPZ, unpublished) reveals key residues (blue) involved in peptide binding.

It has been established that the Nedd4 ligases recognize substrates primarily through their WW domains, small structural domains characterized by a three-strand, anti-parallel β-sheet with two conserved tryptophan residues ~20 amino acids apart (**Figure 1B**).^8,26–31^ WW domains are found in a variety of proteins and bind primarily to proline-rich regions of target proteins. In the Nedd4 family, WW domain-mediated substrate recognition occurs via binding of the WW domain to a substrate PY motif (PPxY or LPxY, where x can be any amino acid; **Figure 1C**), or phospho-threonine or phospho-serine residues (pT and pS, respectively).^26–28,30–36^ There have been various efforts to characterize the nature of Nedd4 interactions with its substrates, from solution state NMR^26,30,33^ and x-ray crystallography^29^ characterization of WW domain-PY motif interactions to pull-down assays and high-throughput microarray screens^34^ of Nedd4 binders. Through these efforts, significant information about the interactions of Nedd4 family has become available with 26 to over 700 interactors annotated for different members of the ligase family (**Table 1**). Despite the availability of this data, there has not been a reported analysis of PY motif-dependent interactions across the Nedd4 family of ubiquitin ligases to date.

**Table 1.** Number of previously identified interactors for each Nedd4 family ligase in the BioGrid protein-protein interaction database.

| **Ligase** | **# of annotated interactors** |
| --- | --- |
| Nedd4 | 348 |
| ITCH | 233 |
| SMURF1 | 425 |
| SMURF2 | 150 |
| WWP1 | 112 |
| WWP2 | 763 |
| HECW1 | 26 |
| HECW2 | 308 |

Using available interactome data, we sought to analyze the features defining the PY-dependent substrates of the ligase family. Through this effort, we aimed to characterize the prevalence of canonical PY motifs (both PPxY and LPxY) in the known Nedd4 family interactome to determine the frequency of PY-mediated substrate recognition. Further, we sought to determine the preferred amino acid identity at the x position and the sequence context of the PPxY and LPxY motifs to provide insight into the nature of the protein regions where these domains occur. To this end, we developed a python-based tool, herein termed PxYFinder, for rapid sequence-based analysis of the Nedd4 family interactome. Analysis of the primary sequence of known Nedd4 family interactors revealed that PY-motifs occur in ~50% of the Nedd4 family interactome, with PPxY motifs occurring more frequently than LPxY motifs. Next, using consensus sequence data from the PY motif-containing interactors of the Nedd4 interactome, we conducted a computational analysis of PY peptide affinity using a rationally designed peptide library. Specifically, we screened combinations of the most common residues at the x_–1_ and x position (where x_–1_ denotes the residue immediately preceding PPxY or LPxY) using a combination of template-based peptide docking^37^ and molecular mechanics-based binding affinity prediction^38^ to identify residue-dependent trends in peptide binding affinity. Finally, to gain insight into PY-independent Nedd4/substrate interactions, we conducted an analysis of the non-PY motif containing Nedd4 substrates to identify possible alternative modes of interaction with the ligase. To this end, we screened non-PY substrates against the PhosphoSite database^39^ to identify phospho-proteoforms that may drive Nedd4 recognition. Cumulatively, the results presented herein provide insight into the predominance and nature of PY-motif dependent protein-protein interactions versus PY-independent substrates in the Nedd4 family interactome and establishes a platform for further experimental interrogation of specificity and substrate affinity in the Nedd4 family of ubiquitin ligases.

## Results

### Identification and analysis PY motif sequences in the Nedd4 family interactome

To begin our analysis of PY motif-mediated interactions in the Nedd4 family, we first sought to determine the prevalence of PY motifs amongst interactors of the family. To this end, we retrieved interactome data for the Nedd4 family ligases (Nedd4–1, Nedd4-2, ITCH, WWP1, WWP2, SMURF1, SMURF2, HECW1, HECW2) from BioGrid^40,41^ using *Homo sapiens* as an organismal filter. There are 26 to 763 annotated interactors for each of the Nedd4 family ligases annotated in the BioGrid database (**Table 1**), so we sought a rapid method to screen the interactor sequences for the presence or absence of PY motifs. While there are numerous bioinformatic tools for identification of characteristic protein domains,^42,43^ there is not, to our knowledge, a tool for rapid identification of PY motifs. Since PY motifs can be identified from the protein primary sequence and do not rely on predicted or annotated protein secondary structure or conformation, we developed a python-based script for rapid analysis of protein primary sequence to identify PY motifs (**Figure 2A**). This script, herein referred to as PxYFinder, and associated documentation are available as supplementary material.

**Figure 2.**
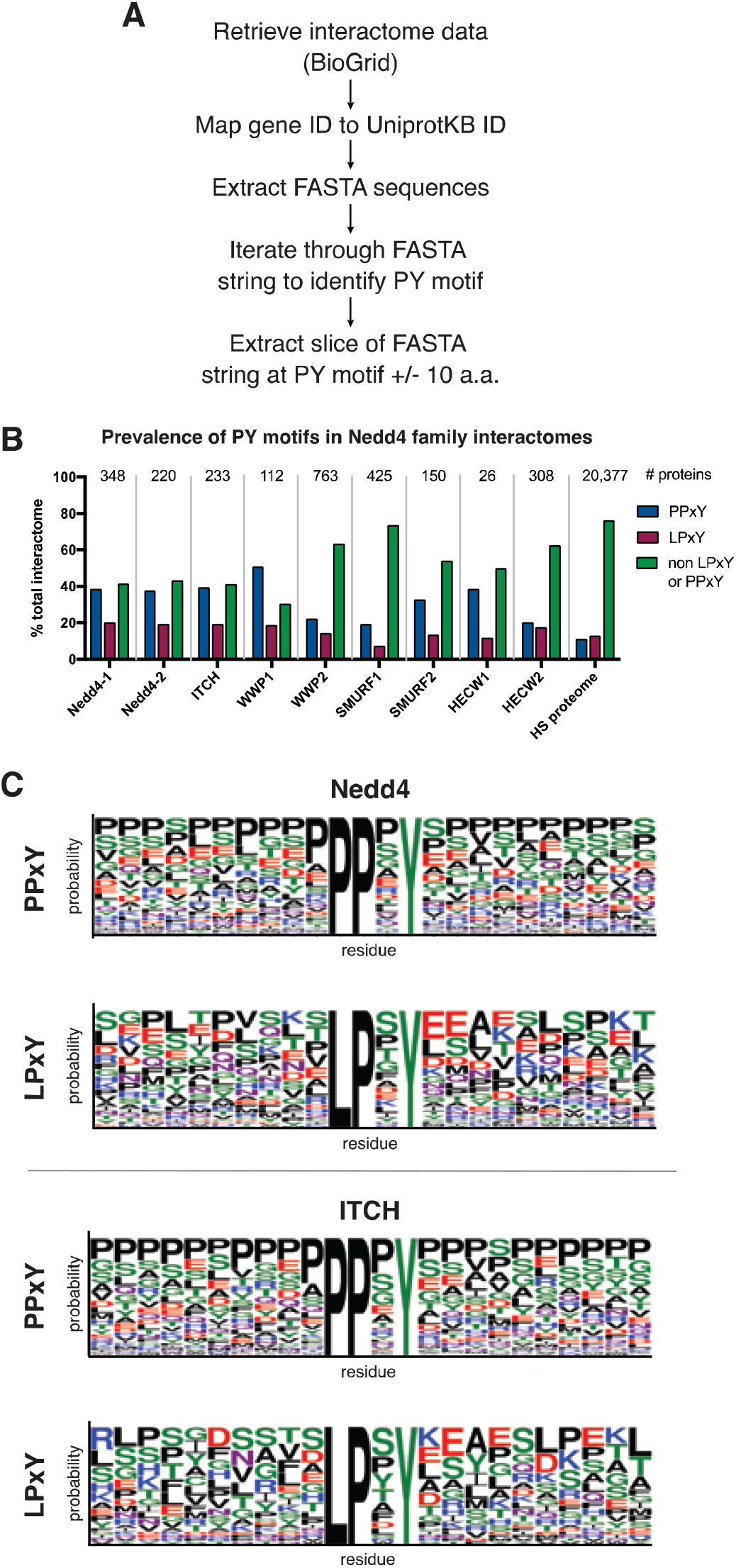
Analysis of PY motifs in interactomes of the Nedd4 family of ubiquitin ligases. **(A)** Workflow for accession of Nedd4 family interactome data and PxYFinder analysis. **(B)** Prevalence of PPxY and LPxY motifs in the interactome (from BioGrid database)^40,41^ across the Nedd4 family of ubiquitin ligases. **(C)** Representative WebLogo depictions of PY motif consensus sequences from all PPxY and LPxY motifs ± 10 amino acids for Nedd4 and ITCH. The WebLogo^44^ diagrams for the remainder of the Nedd4 family ligases are shown in Figure S2.

Using PxYFinder, we identified the prevalence of PPxY and LPxY motifs in the previously annotated interactors of the ligases (**Figure 2B**). To this end, we found that that, on average 33.3% of Nedd4 family interactors contain PPxY motifs while 15.7% contain LPxY and 51.0% contain neither PPxY or LPxY motifs. The prevalence of PY motifs in the Nedd4 family interactomes is enriched relative to the annotated *Homo sapiens* proteome. Our analysis revealed that for all ligases studied, PPxY motifs were more prevalent in the interactome than LPxY motifs. Interestingly, Nedd4–1, Nedd4-2, and ITCH showed similar trends wherein their interactomes were distributed with approximately 40% containing PPxY motifs, 20% containing LPxY motifs, and 40% containing no canonical PY motif. WWP1 showed similar trends with its distribution skewed slightly toward PPxY motif prevalence. For WWP2, SMURF1, SMURF2, HECW1 and HECW2, however, 50% or greater of the interactome lack canonical PY motifs. These results show that on average, approximately 50% of the known Nedd4 family interactome contains a canonical PY motif, providing a sequence based evidence of the likely mode of interaction between Nedd4 and these substrates.

To further understand PY-dependent substrate recognition in the Nedd4 family interactome, we sought to characterize sequences of identified PY motifs to determine 1) if there was conservation of amino acid identity at the x position in the motif and 2) if there are characteristic features of the protein sequences up and downstream of the PY motif. To this end, we used PxYFinder to extract a slice of the FASTA string of each PY motif-containing interactor that included the identified motif and the 10 amino acids before and after the PY motif. After collecting these extracted sequences across the interactome, we looked for consensus sequences using the WebLogo tool^44^ (**Figure 2C, Figure S1**). We see moderate conservation of residue identity at the x position of the PPxY motif across the Nedd4 family, with all but SMURF2, HECW1 and HECW2 having proline, serine, and glycine as the three highest probability amino acids in the x position in the consensus sequences. For LPxY containing proteins, the highest probability residue for the x position is shown to be serine or proline for all members of the Nedd4 family except for SMURF1, HECW1 and HECW2. Interestingly, the WebLogo analysis indicates that PPxY motifs are more likely to occur in proline rich regions of the substrate protein than LPxY motifs. In fact, proline is the highest probability residue at almost all of the up and downstream positions for PPxY motifs in substrates of Nedd4–1, ITCH, WWP1, WWP2, SMURF2 and HECW2. On the other hand, there is little sequence consensus up- and downstream of LPxY across the Nedd4 family, with a distribution of charged, polar, and non-polar residues present as highest probability residues across the consensus sequences.

To gain further insight into the structural context of PY motifs in the Nedd4 family interactomes, we sought to characterize the relative order of the protein regions in which PY motifs occur. Using prototypical member Nedd4 as a case study, we extracted PY motif sequences ± 20 amino acids up and downstream from the FASTA sequences of Nedd4 interactors. We then employed the IUPred2A^45^ tool to calculate the relative order of 109 and 41 sequences containing PPxY and LPxY motifs, respectively. Comparison of the average relative order (wherein a score > 0.5 indicates disorder) of these sequences reveals that, on average across the Nedd4 interactome, PPxY motifs occur in a more disordered region that LPxY motifs (**Figure 3A**). We anticipate that this is a result of PPxY motifs occurring in more proline-rich regions than LPxY motifs (**Figure 2C; Figure S1**) as proline-rich regions are associated with intrinsic disorder due to the geometric constraints imposed by the backbone of proline residues.^46,47^ To further analyze the effect of proline prevalence on the predicted order of the PY motif-containing regions, we employed PPIIPred,^48^ a bioinformatic tool for identification of polyproline II (PPII) secondary structure, an extended helix-like structure that can occur in the presence of polyproline stretches. PPIIPred analysis reveals that PPxY motifs are more likely to display PPII structure immediately before or at the PY motif (residues 20-24 in extracted sequence slices) as compared to LPxY motifs (**Figure 3B**). The sequences up and downstream of the PY motifs, however, show similarly low propensities for PPII structure on average. We anticipate that the increased prevalence of proline residues in PPxY-containing regions contributes to relative disorder but does not induce PPII structure on average.

**Figure 3.**
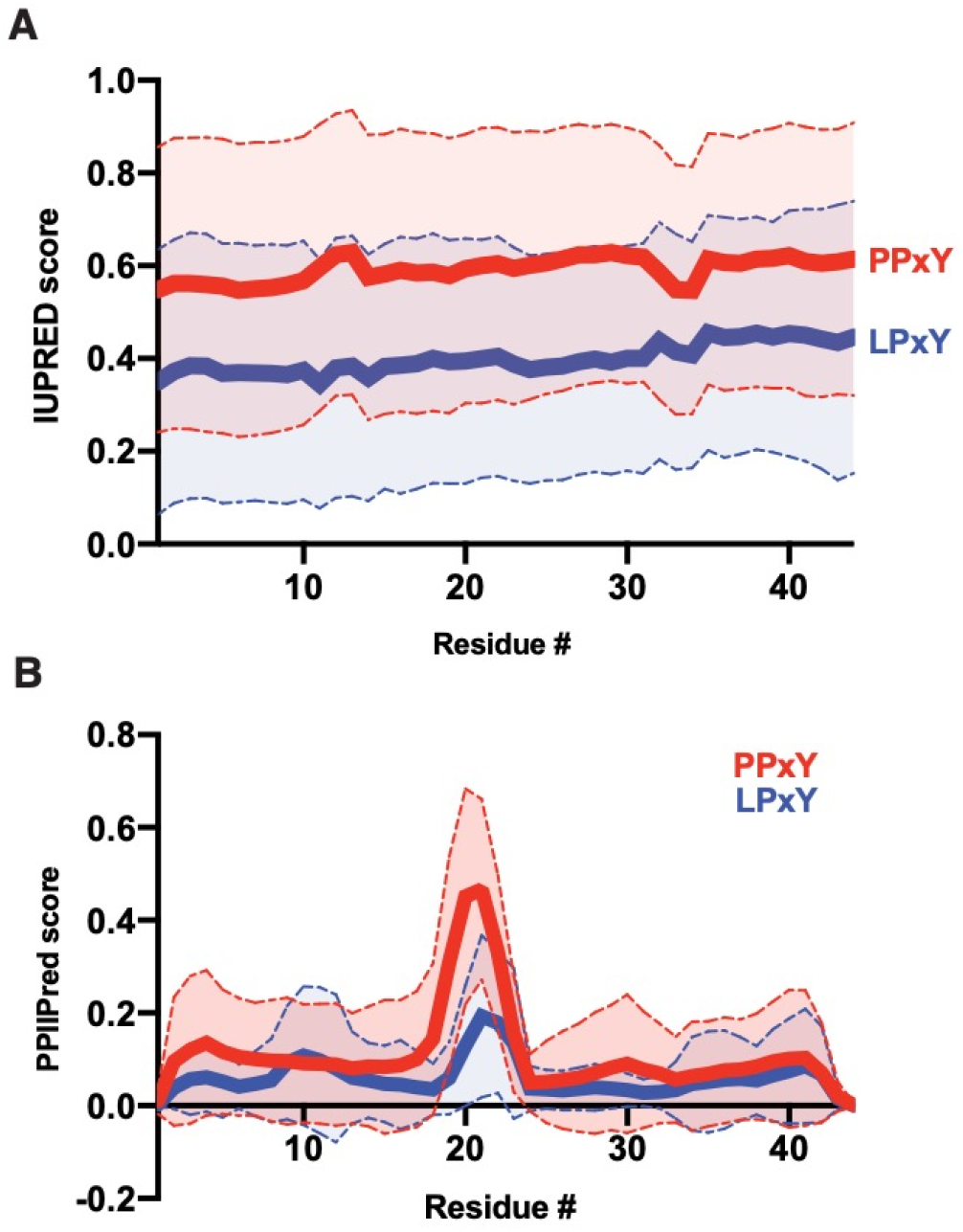
Prediction of relative order of PY-motif containing regions from the Nedd4 interactome. PY motif sequences ± 20 amino acids were extracted from the Nedd4 interactome using PxYFinder script and analyzed using **(A)** IUPred2A^45^ and **(B)** PPIIPred^48^ bioinformatic tools to determine relative order and propensity to form polyproline II structure, respectively. Data shown as average ± S.D. of 109 (PPxY) and 41 (LPxY) sequences. Statistical analysis using a paired t-test (to compare at each residue, numbered 1–44 above) reveals a statistically significant difference in predicted order between the PPxY and LPxY sequences (p < 0.0001 for both IUPred and PPIIPred scores). Analysis across the sequence using an unpaired Welch’s t-test also shows significant differences (p < 0.0001 for IUPred; p < 0.002 for PPIIPred).

### Rational design of PY motif peptide library for computational analysis

Analysis of previously resolved PY motif/WW domain complex structures from Nedd4 show moderate conservation of PY peptide backbone conformation regardless of primary sequence (**Figure 4A**). To better understand the effect of PY sequence on WW domain binding, we sought to determine if the sequence variants of the PY motif affected the predicted affinity with which the substrate of interest binds to Nedd4. To this end, we began with a previously resolved structure of a Nedd4 WW domain bound to a PY-motif peptide from a known substrate, sodium channel ENaC (PDB ID: 2M3O) as our model complex.^30^ Using the ENaC peptide (sequence: TAP**PPAY**ATLG, with PY motif in bold) as a template, we designed a peptide library based on the previously described consensus sequences. We chose to vary the residues at the x and x_−1_ position of the PY motifs (where x_−1_ is the residue immediately preceding PPxY or LPxY) as these are the residues which span the binding interface between the PY peptide and WW domain (**Figure S2**). Based on the consensus sequences (**Figures 2C, S1**), we generated 15 variants each for PPxY and LPxY peptides using the template peptide (TA**x**_**−1**_**PPxY** ATLG or TA**x**_**−1**_**LPxY** ATLG) with all combinations of the three and five highest probability residues at the x_−1_ and x positions, respectively (**Figure 4C**). It should be noted that, as there is not a previously characterized complex of a hNedd4 WW domain bound to a PY-peptide with the LPxY motif, we opted to use the same template peptide for screening of both PPxY and LPxY motifs to allow for direct comparison across the suite without variation outside of the peptide core. Cumulatively, this design afforded 30 peptides in total for computational screening against a Nedd4 WW domain.

**Figure 4.**
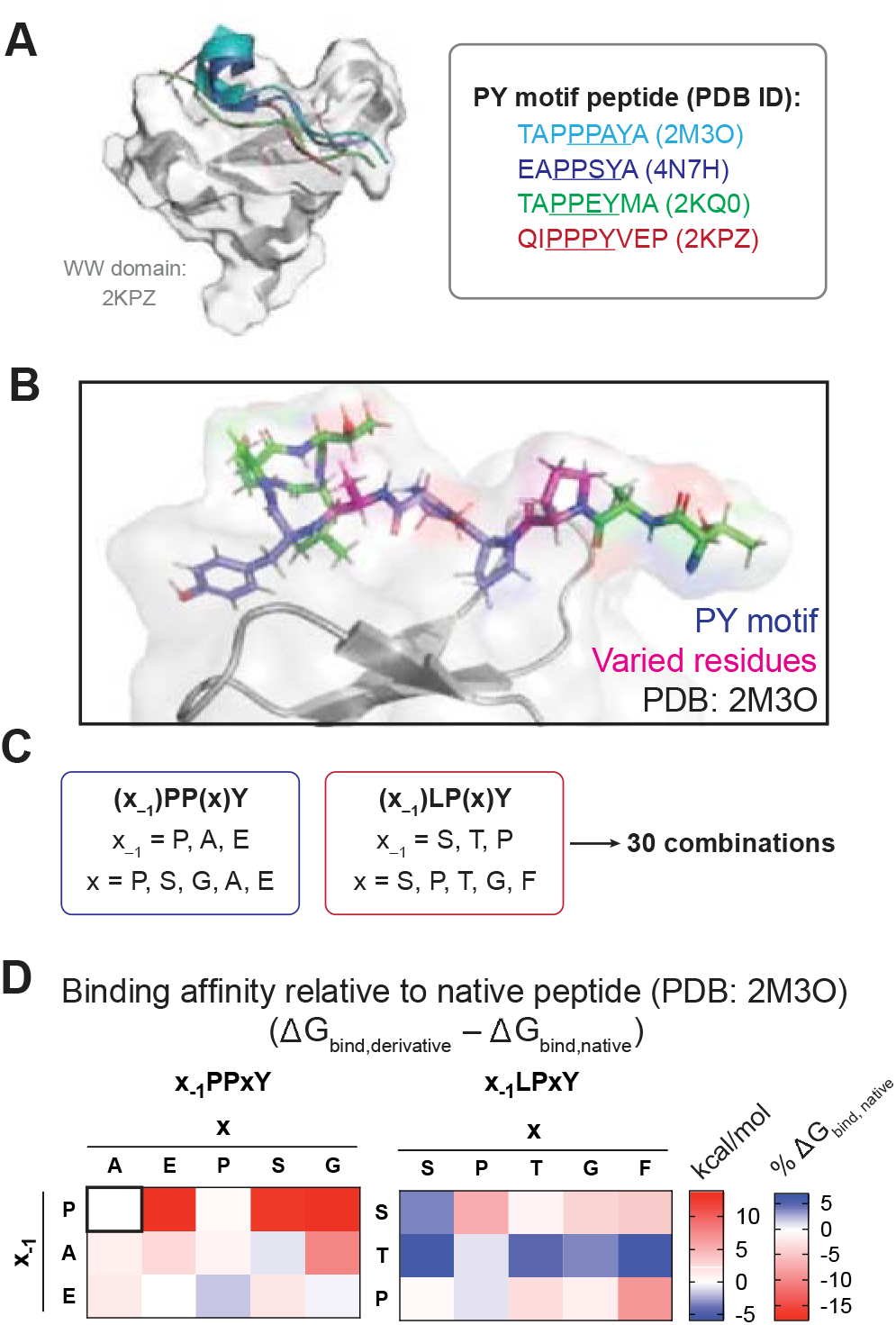
Rational design and computational analysis of PY motif peptide library to predict residue-specific changes in binding affinity. **(A)** Nedd4 WW domain (PDB ID: 2KPZ) in complex with PY motif peptides from previously resolved WW domain/PY peptide complexes (PDB IDs: 2KPZ, 2M3O, 2KQ0, 4N7H). Peptides aligned to the 2KPZ complex using PyMol,^58^ showing moderate conservation of peptide backbone conformation when bound to the WW domain. **(B,C)** Rational design of PY peptide library involved variation of residues in the x_−1_ and x positions (shown in pink, **B**) and was informed by PY motif consensus sequences for Nedd4 (shown in Figures 2C and S2), affording a 30 member library. **(C)** Computationally predicted binding affinities of PY motif peptides screened against Nedd4 WW domain (PDB ID: 2M3O). Binding affinities are presented as ΔΔG_binding_ relative to the native peptide substrate TAPPPAYATLG (ΔG_binding_^designed^ – ΔG_binding_^native^). ΔΔG for the native peptide (PPPAY) is presented in the upper-right corner of the left heatmap for reference. ΔΔG_binding_ energies presented as kcal/mol or % of ΔG_binding_^native^ where blue and red indicate a more and less favorable (negative) binding energy, respectively. Full energy properties described in Table S2 and as .csv in Supplemental Material.

### Docking and molecular mechanics analysis of WW domain/PY motif interactions

Prior to computational analysis of our rationally designed peptide library, we first sought to determine the WW domain scope required to capture any sequence-dependent variation in WW domain binding across the Nedd4 family. To this end, we compared the conservation of WW domain sequences across the family of ligases. Each ligase contains 2–4 WW domains (**Figure 1A**), with moderate sequence similarity across the family (**Figure 1B, Figure S2A,B**). Analysis of key residues that interact with peptide substrates shows moderate conservation of the binding interface (**Figure S2C**). These residues, which are primarily located in the concave peptide binding cleft of the three-stranded β-sheet structure, drive the direct interaction of the WW domain with the PY motif (**Figure S2C**). Alignment of three representative WW domain structures with varied residue identity in the binding interface (Nedd4–1 (WW3) and ITCH (WW3 and WW4), all of which have been shown to bind ENaC, the substrate from which the peptide library was derived) shows conservation of secondary structure (**Figure S2D**). Additionally, analysis with MolProbity^49^ and KiNG indicate^50^ that the interactions between the peptide substrate and WW domain are predominantly mediated by van der Waals contacts with few inter-peptide and peptide-WW domain hydrogen bonds (**Figure S2C**). Therefore, we anticipate that trends in binding affinity across the peptide library from screening against our model WW domain (PDB: 2M3O) will be representative as the electrostatic nature of the binding interface is largely conserved. Instead, we anticipate that sequence-dependent changes in the peptide-protein interactions and peptide conformation will have a greater effect on binding affinity than the identity of WW domain residues.

To begin our computational analysis of the rationally designed PY motif library against our suite of WW domain structures, we first sought a docking method that was amenable to docking peptides to protein targets rather than small molecule ligands. We determined that use of a template-based docking method was the most appropriate approach for our analysis as there are a number of PY peptide/WW domain complexes that have been reported. Therefore, known PY/WW complex structures can be used to guide the docking, improving efficiency and minimizing computational expense compared to a global docking approach. To this end, we employed GalaxyPepDock^37^ to dock our library against a Nedd4 WW domain (PDB ID: 2M3O).^30^ Each GalaxyPepDock docking result provided 10 predicted poses for the peptide of interest. From these 10 poses, we selected the pose with the most similarity to the native substrate in the PY/WW complex (PDB ID: 2M3O) with particular emphasis on the position of the tyrosine residue in the PY motif, which forms key dipole-ion interactions with conserved lysine and histidine residues in the WW domain (**Figures S2 and S3**). To further refine the docking result, we then used the Glide ligand docking tool with SP-Peptide precision setting from the Schrödinger suite^51–53^ to optimize the conformation of the peptide backbone and side chains in the binding pocket. Finally, to obtain thermodynamic measurement of predicted peptide binding affinities, we employed molecular mechanics-based binding affinity prediction using the Generalized Born and surface area continuum solvation method (MM/GBSA, Schrödinger),^38^ which considers the effect of solvation on binding energies using an implicit solvation model. From this calculation, we generated a total measurement of affinity as ΔG_binding_ in addition to various contributing energy terms, enabling analysis of biomolecular interactions that serve as driving forces in the peptide/protein interaction.

The predicted ΔG_binding_ values provide a relative measure of affinity across the peptide suite wherein a more negative number indicates a stronger predicted peptide/protein interaction. Docking results were analyzed by comparison to the predicted binding affinity of the native peptide (**Figure 4D**). Our docking analysis reveals that, in general, substitutions at the x_−1_ or x position in the PPxY peptide scaffold weaken the predicted binding affinity (indicated by a less negative ΔG_binding_) with the exception of derivatives APPSY, EPPPY, and EPPGY. We anticipate that there is a significant deal of pre-organization in the native ligand around the tri-proline core (TA**PPP**AYATLG), and we hypothesize that alteration of the steric or electrostatic nature at the x position with retention of the tri-peptide core (PPPxY) is unfavorable as the peptide lacks flexibility to compensate for altered interactions with the WW domain. Screening of derivatives with alanine or glutamic acid at the x_−1_ position were slightly unfavorable, but derivatives APPSY, EPPPY demonstrate improved affinity, likely through an increased number of intramolecular interactions due to the bent conformation adopted by the optimized docked ligand (**Figure S4**).

In the LPxY peptide library, the derivatives generally had stronger predicted binding affinities than the PPxY library members. We anticipate that this is a result of greater ligand flexibility resulting from the lessened conformational strain induced by the core proline-proline dipeptide. Several members of this peptide class that have strong predicted binding affinities adopted a bent conformation, increasing the number of intramolecular contacts. Further, we hypothesize that the increased polarity with substitutions of serine or threonine at the x_−1_ or x positions increases either dipole-mediated intramolecular interactions or stabilizes the peptide/WW domain complex by presenting the polar residue to the solvent accessible side of the peptide and promoting burial of the lipophilic residues in the WW domain binding pocket.

We then analyzed individual energetic contributions to overall binding affinity across the library of peptide analogues. This includes energy components of the free ligand or receptor, the optimized complex, or sub-components of the ΔG_binding_ measurements (i.e. contributions of individual interaction types). In general, van der Waals and Coulombic interactions contributed most strongly to binding affinity, while solvation energy accounted for the most disfavorable (positive ΔG) component (**Figure 5A and S5**). We next correlated all individual energy components to total ΔG_binding_ (**Figures 5B,C and S6**). Analysis of energy components from complex, ligand, and receptor showed that receptor energies had the lowest correlation with overall ΔG_binding_ while ligand and complex energy components had higher correlations (**Figure 5B**). Further analysis showed that ligand efficiency, a function of binding affinity relative to total non-hydrogen atoms, correlated most strongly with ΔG_binding_ (**Figure 5C**). Finally, this analysis reveals that van der Waals are most correlated with ΔG_binding_, followed by Coulombic and lipophilic interactions.

**Figure 5.**
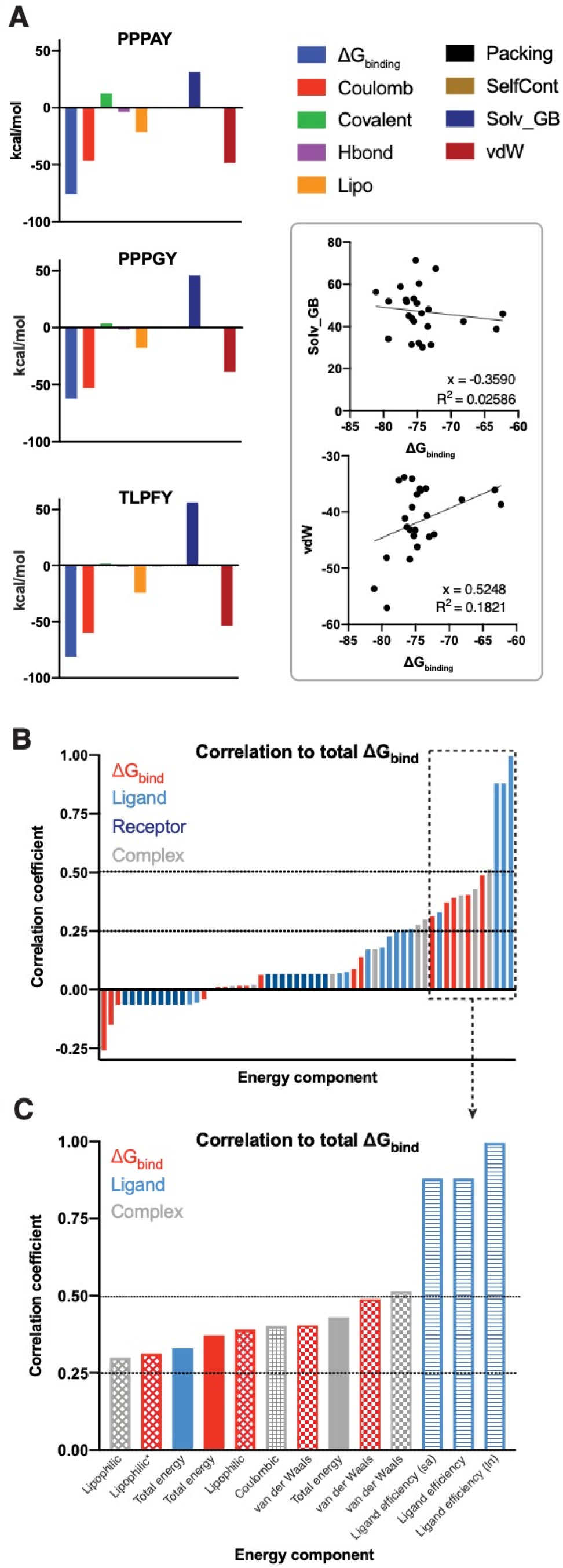
Individual biomolecular interaction types have varying contributions to overall binding affinity. **(A)** Energetic contributions of individual interactions to overall binding affinity are shown for the native ligand (PPPAY), a weaker predicate binder (PPPGY), and a stronger predicted binder (TLPFY). Linear regression analysis reveals a positive and negative correlation, respectively, between van der Waals forces or solvation energy with overall binding energy (ΔG_binding_). **(B)** Analysis of all individual energy components (for ΔG_bind_, the optimized complex, ligand, and receptor) to overall binding affinity reveals factors that ligand and complex energies are more strongly correlated than receptor energies. Ligand efficiency is defined as the binding energy/# heavy atoms where “sa” accounts for solvent exposed surface area and ln is the natural log of ligand efficiency. * indicates lipophilic interactions in the ΔG_bind,NS_ where NS indicates binding energy of the peptide without accounting for ligand strain energies. Correlations calculated using python as Pearson coefficients and visualized in Prism GraphPad.

### Analysis of non-PY motif substrates in the Nedd4 interactome

While we have extensively discussed the nature of PY motif-mediated interactions with WW domains, the nature of interactions that guide the remaining half of the interactome remain unclear from our analysis. It is likely that these interactions are guided by interactions at other sites in the ubiquitin ligase, such as is the case for E2 conjugating enzymes,^54–56^ which interact with the HECT domain, or for proteins like α-synuclein,^11–14,16^ which has been shown to interact with the C2 domain and HECT domain of the ligase. Additionally, there is evidence for WW domain interactions with phospho-threonine or phospho-serine (pT, pS).^26^ In these cases, the Nedd4 interaction would be dependent upon specific phospho-proteoforms, the presence of which are regulated by other cellular pathways and is discussed further below.

To complement our analysis of PY motif-mediated protein-protein interactions in the Nedd4 family interactome, we sought to further analyze the pool of non-PY motif interactors in the annotated dataset. We first performed a functional analysis of non-PY interactors using the PANTHER Gene Ontology annotation database^57,58^ to determine how many proteins in the interactome were involved in the ubiquitination process (for example, E2 conjugating enzymes that would bind to the ligases through the E2 interaction site on the catalytic HECT domain). From this annotation, we identified that the non-PY containing interactome contained a range from 2.19% (WWP2) to 11.63% (WWP1) across the Nedd4 family of Nedd4 (**Table 2**). This suggests that nearly all of the non-PY interactome is not comprised of upstream members of the ubiquitin signaling cascade but rather contains substrates or regulatory partners that interacts with Nedd4 in a PY-independent manner (i.e. through phosphorylated residues or through C2 or HECT domain interactions).

**Table 2.** Functional analysis of non-PY containing interactors involved in ubiquitination.

| **Ligase** | **# non-PY motif interactors** | **% of non-PY interactome involved in ubiquitination** |
| --- | --- | --- |
| Nedd4 | 144 | 3.73 |
| ITCH | 96 | 11.46 |
| SMURF1 | 312 | 5.36 |
| SMURF2 | 81 | 10.94 |
| WWP1 | 34 | 11.63 |
| WWP2 | 483 | 2.19 |
| HECW1 | 14 | 42.8 |
| HECW2 | 213 | 4.25% |

To characterize the remainder of the non-PY containing proteins in the Nedd4 interactome, we next screened non-PY interactors for the presence of pT or pS residues as reported in the PhosphoSite protein phosphorylation database. Of the 153 non-PY interactors identified in the Nedd4 interactome, there are 128 proteins that are annotated in PhosphoSite database^39^ to contain both pT and pS post-translational modifications (PTMs) while 17 proteins have been detected with either pT or pS and eight proteins have no reported pT or pS residues. Based on these previous reports, experimentally detected phosphorylation on threonine and/or serine residues occurs in 94.8% of the non-PY interactome. Thus, this provides putative evidence that phosphorylation at serine or threonine may be the driving force for Nedd4 recognition of non-PY substrates. Though experimental validation of putative WW domain/phospho-protein interaction specificity would be required, it is beyond the scope of this investigation.

## Discussion

We have employed a combination of bioinformatic and computational analyses to gain insight into the sequence and structural properties that drive interaction specificity in the Nedd4 interactome. We began our analysis with the development and implementation of PxYFinder to rapidly identify the presence or absence of canonical PY motifs in a library of FASTA sequences. Using this tool in combination with interactome data available through the BioGrid database, we determined that, on average, 33.3% of Nedd4 family interactors contain PPxY motifs while 15.7% contain LPxY and 51.0% contain neither PPxY or LPxY motifs. This demonstrates that canonical PY motifs drive only half of the WW-domain mediated interactions in the known interactome on average, and that screening for PY motifs is not sufficient on its own for identification of putative Nedd4 family substrates. In general, all members of the Nedd4 family that we analyzed have more interactors that contain PPxY motifs than LPxY motifs, and consensus sequence analysis reveals that PPxY motifs are more likely to occur in proline rich regions than LPxY motifs.

Using the information obtained from PxYFinder and consensus sequence analysis of the Nedd4 interactome, we sought to computationally analyze sequence-dependent effects on PY peptide/WW domain binding. To this end, we employed a multi-step computational analysis of peptide binding affinity using a previously resolved structure of the Nedd4 WW domain. Specifically, we designed a library of PY peptides (both PPxY and LPxY motifs) which contain all combinations of the three and five most commonly occurring residues at the x_−1_ and x positions based on our bioinformatics-derived consensus sequences. We then employed a multi-step computational analysis wherein we began with template-based docking of the peptide substrate to the WW domain structure, followed by refinement of the complex and analysis of thermodynamic binding parameters via MM-GBSA. From this effort, we determined that the PPxY scaffold is less tolerant to substitutions than the LPxY scaffold. We hypothesize that this is a result of pre-organization and strain in the poly-proline backbone of the PPxY peptides. Therefore, incorporation of residues that increase peptide flexibility or polarity tend to improve binding affinity. As predicted, our analysis reveals that binding affinity is most strongly driven by van der Waals interactions, with positive though lesser correlations to Coulombic and lipophilic interactions.

To gain further insight into the role of WW-domain binding in Nedd4 family substrate recognition, we analyzed the non-PY containing interactome of Nedd4 as a case study. This analysis reveals that nearly all of the non-PY substrates have been previously annotated to have phosphorylation at threonine and/or serine residues, providing a putative indication of WW-domain recognition independent of canonical PY motifs. While experimental validation of these hypotheses would be necessary to confirm the mechanism of Nedd4 recognition, our bioinformatic analysis provides valuable insight into possible modes of binding.

Cumulatively, the results presented herein provide insight into the prevalence and nature of PY motifs in the Nedd4 interactome. We anticipate that PxYFinder will be useful in screening large datasets for putative WW-domain interactors (both in the Nedd4 family and for other WW domain-containing proteins) and addresses a gap in current bioinformatic tools for which there is not an established method for identification of PY motifs in a large dataset. Further, our analysis of identified PY motifs expanded our understanding of the conservation of residues in and around the motif, and informed a computational analysis of sequence-dependent changes in PY peptide binding affinity. While the binding parameters obtained in this computational analysis are relative, we anticipate that our results will be useful in informing experimental design of PY peptide libraries either for interrogating the nature of the peptide/protein interaction or for designing inhibitors that target PY peptide/WW domain complexes.

## Methods

### Development and use of PxYFinder script

A python script, termed PxYFinder, was developed in Python 3.8 to perform the following workflow: PxYFinder imports FASTA sequences and iterates through primary sequences to identify PPY, PPxY, or LPxY. If a PY motif is identified, PxYFinder extracts a slice of the FASTA string that contains the PY motif and x (user-specified) amino acids up and down stream, copying this slice to a new .csv file. Code and documentation for PxYFinder is available in the Supplementary Material.

For analysis of Nedd4 family interactors, interactome data for each Nedd4 family member of interest (Nedd4, ITCH, WWP1, WWP2, SMURF1, SMURF2, HECW1, HECW2) was retrieved from BioGrid using *Homo sapiens* as an organismal filter. Gene names were converted to UniprotIDs and were used to retrieve FASTA sequences from the Uniprot database. PxYFinder was used to identify and extract PY motifs in the interactome of each ligase and calculate prevalence of PY motifs in each interactome. Graphs were generated using Prism (GraphPad). PY motif consensus sequences were determined by analysis with WebLogo (http://weblogo.threeplusone.com/)^44^ with probability as the y-axis measure.

### Prediction of protein order in PY-motif containing segments

PxYFinder script was modified to extract PY motifs ± 20 amino acids from Nedd4 interactor sequences. These slices were subsequently analyzed using the IUPred2A and PPIIPred tools for prediction of overall disorder (IUPred2A)^45^ and propensity to form polyproline secondary structures (PPIIPred).^48^ Data was visualized as mean ± S.D. and analyzed using paired and unpaired (Welch’s) t-test to determine statistical differences between specific residue positions (numbered 1–44) and across the full sequence, respectively. Data visualization and analysis was performed in Prism (GraphPad).

### Docking and molecular mechanics analysis of PY peptide library

A 30-member peptide library was generated based on the consensus sequences in the PY motif interactome of Nedd4. The top three and five residues at the x_−1_ and x positions respectively were paired in all possible combinations to generate 15 peptides each for PPxY and for LPxY libraries using a previously characterized substrate peptide bound to a Nedd4 WW domain (PDB ID: 2M3O)^30^ as a template. All peptides were first docked via GalaxyPepDock^37^ using 2M3O as a template structure. Docked complexes were further refined using the Schrödinger suite (Schrödinger Release 2020-3, Schrödinger, LLC, New York, NY, 2020). Specifically, the complex generated using GalaxyPepDock was prepared using the Protein Preparation Wizard and LigPrep tools.^57^ Using the Glide tool,^51^ a docking grid for the WW domain was generated using (Glide Receptor Grid Generation), and the ligand was docked as a flexible ligand to the generated grid using SP-Peptide function with retention of amide bond conformation and restriction of docking poses to 0.50 Å tolerance for core pattern comparison relative to the native ligand conformation.

Following docking, the Schrödinger Prime MM-GBSA^38,52,53^ tool was used to analyze the PoseView (PV) docking output file with the VSGB solvation model and OPLSe3 force field to generate ΔG_binding_ for each generated pose. For each peptide, the pose with most negative ΔG_binding_ was selected for comparison of predicted binding affinities across the peptide library. Graphs of binding data were generated using Prism GraphPad. Structural analysis and visualization were performed with PyMol,^58^ MolProbity,^49^ and/or KiNG.^50^

### Analysis of non-PY substrates across the Nedd4 interactome

The PhosphoSite^39^ database was used to screen non-PY substrates of the Nedd4 interactome for the presence of previously reported phospho-threonine or phospho-serine posttranslational modifications. The PANTHER Gene Ontology^59,60^ database was used to screen the non-PY substrates for proteins that are known components of the ubiquitin signaling pathway.

## Acknowledgements

The authors would like to thank the Duke University Department of Chemistry Computing Services for access to the Schrödinger software suite and computational servers. They also thank the department for support for M.D.P. through the summer undergraduate research fellowship program. Finally, the authors thank the members of the McCafferty lab for their thoughtful feedback on the project and manuscript.

## Author contributions

A.K.H. designed the project, participated in data collection, analysis, and visualization, prepared the manuscript, and mentored M.D.P. M.D.P. prepared the PxYFinder python script and contributed to data collection and analysis and in manuscript preparation. D.G.M. mentored A.K.H. and M.D.P., secured funding, oversaw the project, and participated in manuscript preparation.

