## Supplemental Materials for "Bioinformatic and computational analysis reveals the prevalence and nature of PY motif-mediated protein-protein interactions in the Nedd4 family of ubiquitin ligases" (from another version of this preprint: version 1, 2020-11-12)

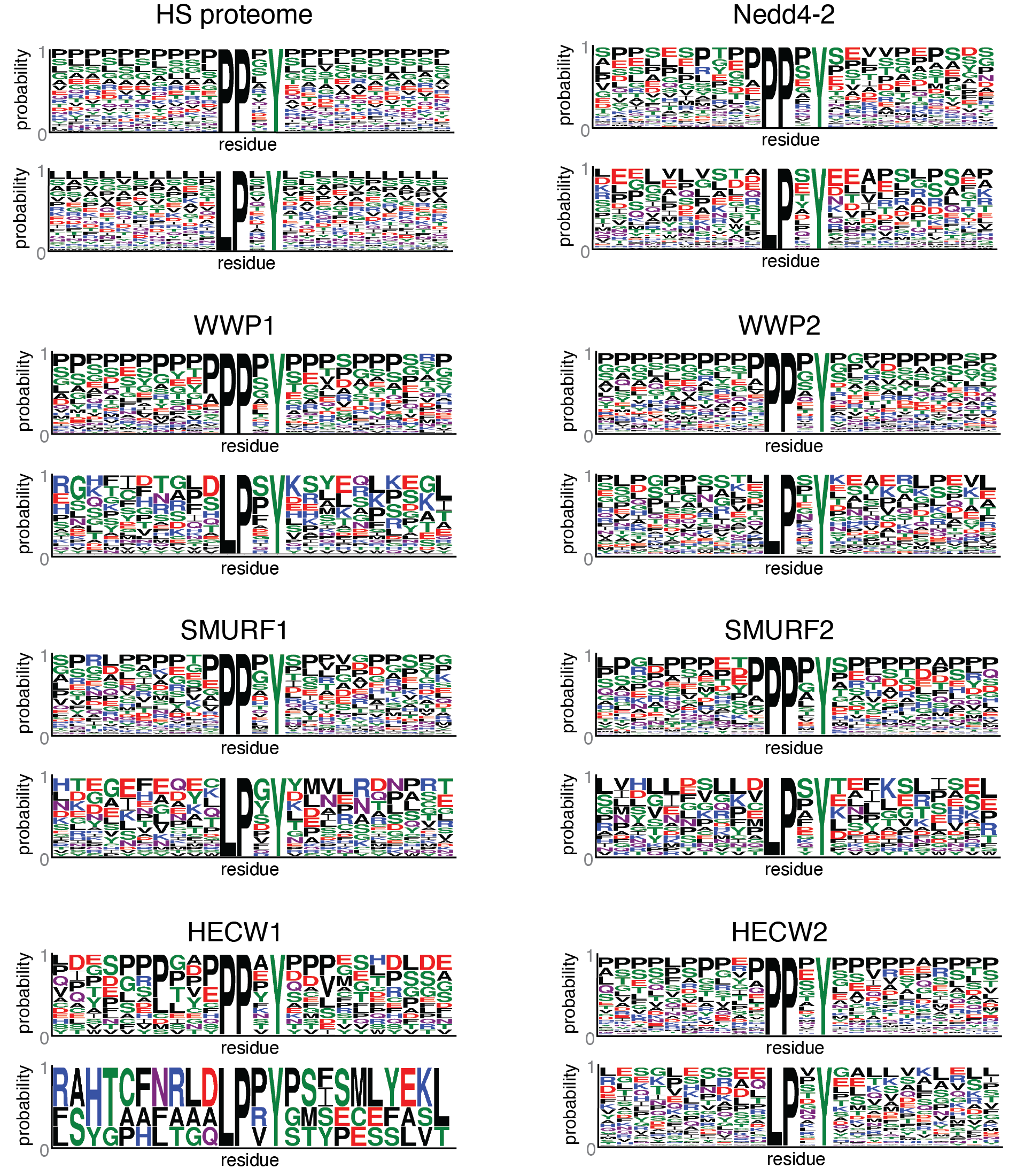
**Figure S1. Sequence Logo analysis reveals trends in PY motif consensus sequences across the Nedd4 family.** Sequence logo diagrams were used to identify consensus sequences in PY motifs and in surrounding regions (± 10 amino acids) for Nedd4 family members and for all *Homo Sapiens* proteins that have SwissProt annotation available in the UniProt database (labeled as HS proteome). Sequence logos for Nedd4-1 and ITCH are excluded from this figure as they are presented as representative images in Figure 2. Sequence logo analysis reveals that PPxY motifs are more likely to occur in proline-rich regions than LPxY motifs, and amino acid identity at the x position is more conserved in PPxY motifs across the Nedd4 family and proteome than in LPxY motifs.

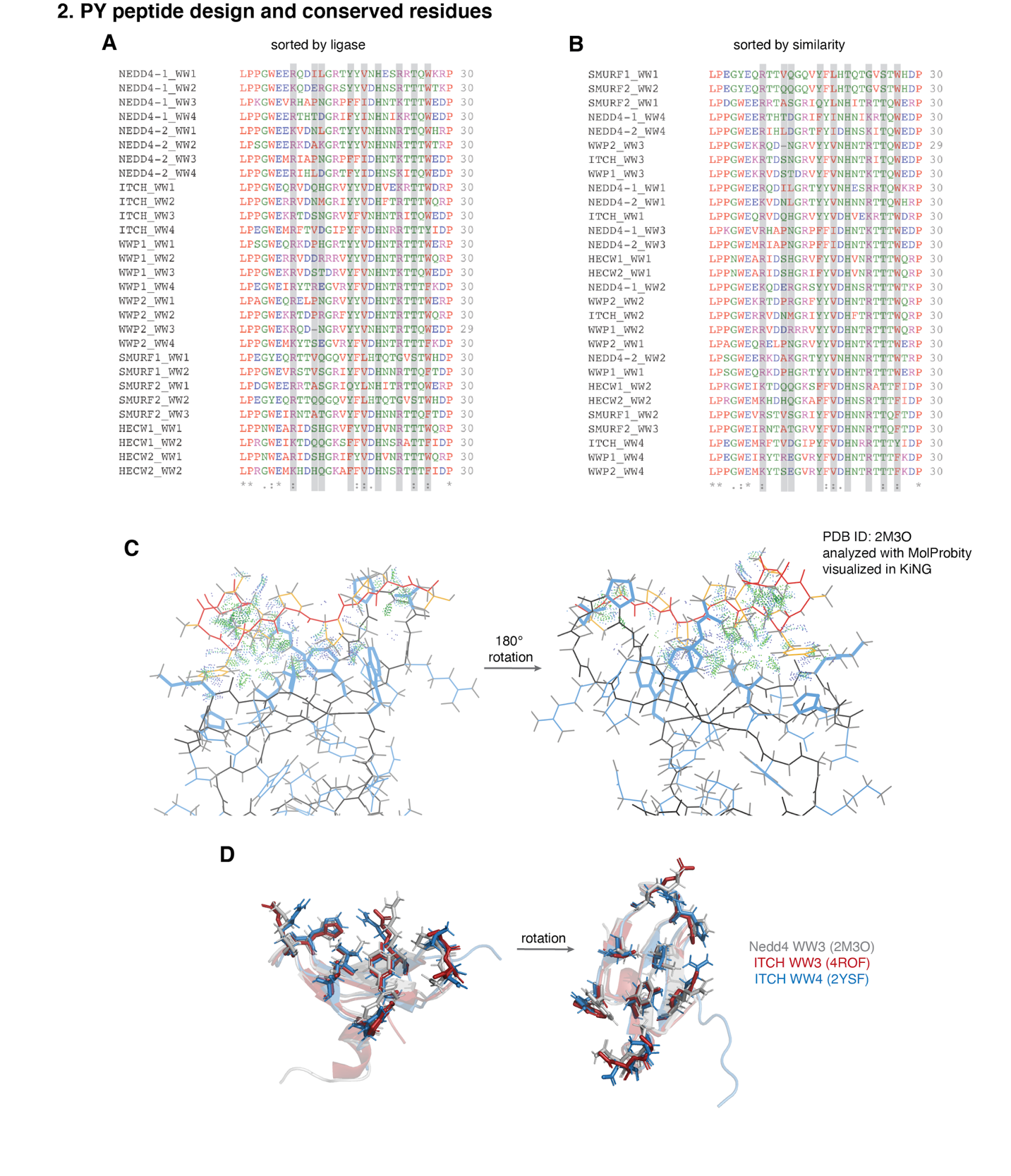

**Figure S2. Nedd4 family WW domain sequence and structure alignment shows moderate sequence and high structural similarity.** Sequence alignments of WW domains from Nedd4 family members sorted by **(A)** ligase and **(B)** similarity show moderate sequence conservation, with high conservation of key residues in the binding interface (highlighted in grey). **(C)** MolProbity analysis reveals contacts between PY peptide substrate (red/yellow) and WW domain binding cleft residues (blue) shows that highly conserved residues (grey in **A** and **B**) are located in the peptide binding cleft. **(C)** Alignment of three WW domain structures with varying sequence similarity show high conservation of structure and of positioning of key residues despite differences in residue identity.

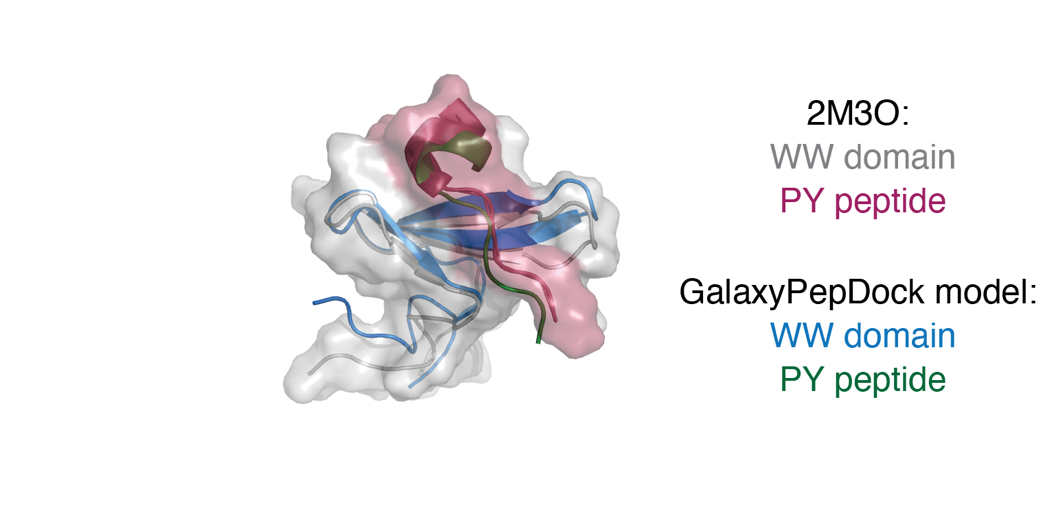

**Figure S3. GalaxyPepDock accurately predicted conformation of substrate peptide based on template.** As a test of GalaxyPepDock template-based docking reliability, the native peptide substrate of Nedd4 WW domain (reported in PDB structure 2M3O) was docked to the apo-WW domain, extracted from PDB 2M3O. Alignment of the native complex (2M3O, peptide shown in red; WW domain in grey) with the docked complex (via GalaxyPepDock; peptide in green; WW domain in blue) show reliable docking of the peptide with retention of conformation and peptide-WW domain contacts.

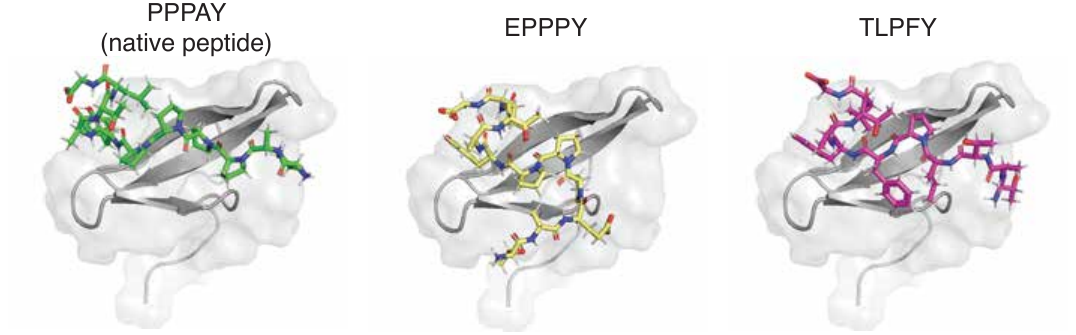

**Figure S4. Conformations of selected peptide derivatives after computational docking and optimization.** A sampling of peptide conformations from computational docking of the rationally designed peptide library demonstrates the variety of intramolecular contacts that the PY peptides can form with the WW domain structure. Binding energies of the representative peptides shown here are presented in Figure 4.

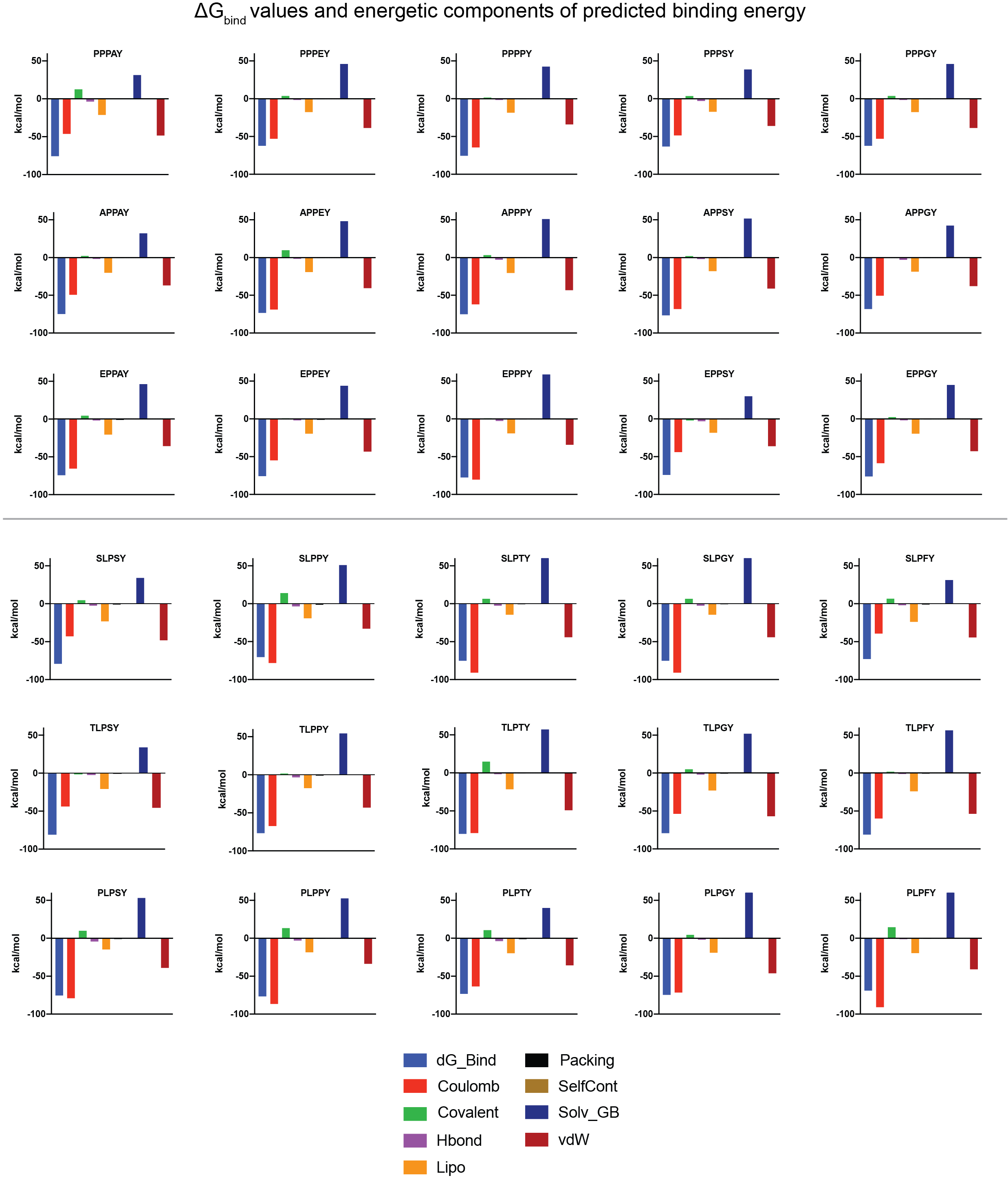

**Figure S5. Energetic contributions to computationally predicted ΔG_binding_ of PY peptide library to Nedd4 WW domain.** ΔG_binding_ and energetic components that contribute to ΔG_binding_ are shown here as calculated with the Schrodinger Prime MM-GBSA tool. Energies are given in kcal/mol, and energy contributions are shown for all 30 members of the rationally designed PY peptide library.

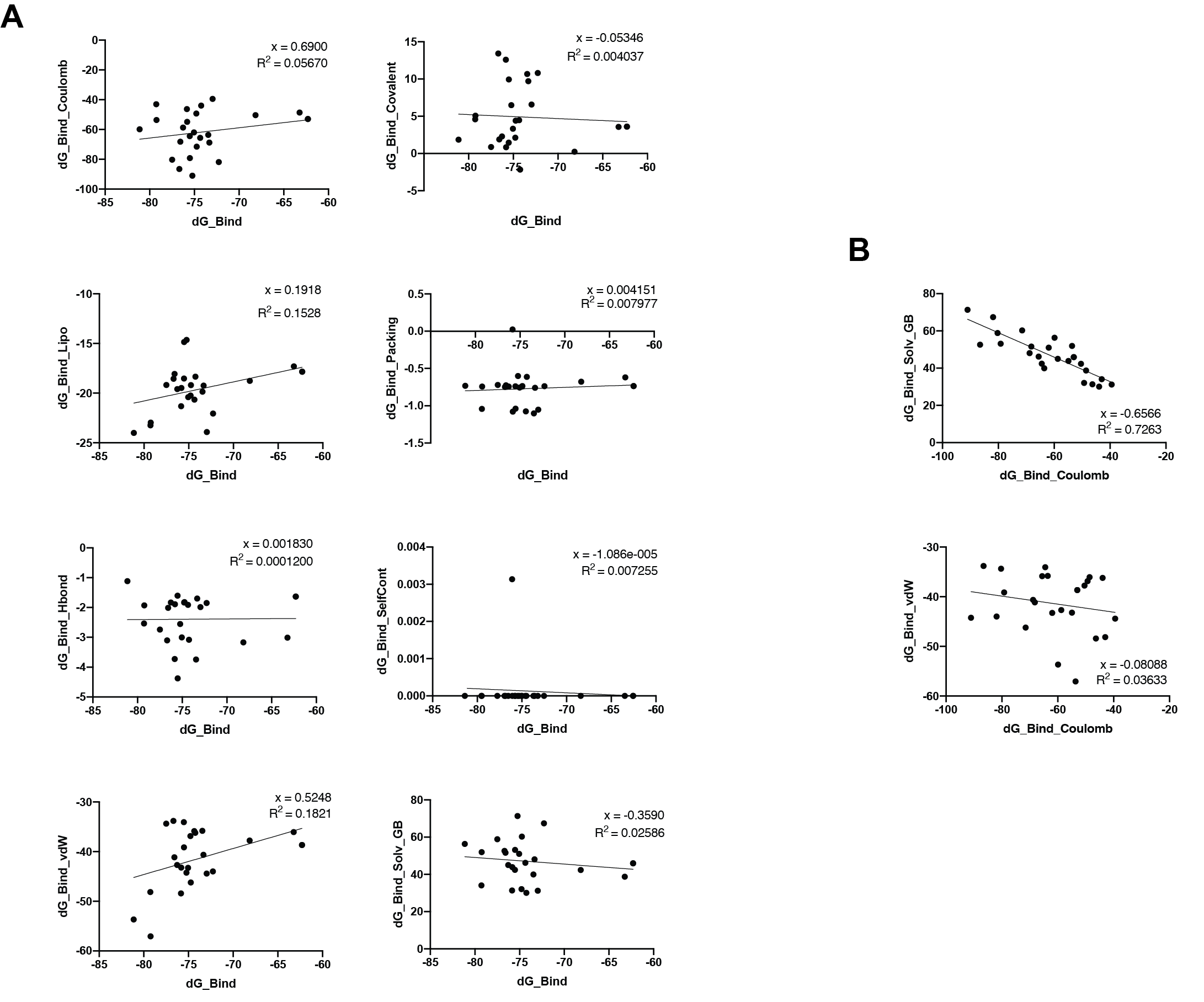

**Figure S6. Correlation of energetic components that contribute to peptide binding.** Correlation of calculated energies (ΔG_binding_ and ΔG_binding_ sub-components) across the peptide library show that **(A)** some energetic contributions are more strongly correlated to overall binding (ΔG_binding_) relative to other components. **(B)** Correlation of solvation (Solv_GB) and van der Waals (vdW) components of ΔG_binding_ with coulombic interactions shows that solvation is more strongly correlated with coulombic interactions than van der Waals interactions Specifically, stronger (more negative) coulombic interactions correlate with more positive solvation energies. Values calculated with Schrodinger Prime MM-GBSA and presented in kcal/mol. Simple linear regression analysis and data visualization performed in Prism GraphPad. X = slope of linear regression line of best fit; R^2^ provided as measure of goodness of fit.
